# Low-cost and highly efficient: A method for high-quality nucleic acid isolation from cotton fibres

**DOI:** 10.1101/2022.10.07.511236

**Authors:** Mukhtar Ahmed, Muhammad Bilal Sarwar, Ramla Ashfaq, Adnan Ahmed, X. Yanang, M. Fanglu, Salah-ud-Din, Muhammad Sajid, Quratulain Syed, Syed Hussain Abidi, Xuede Wang

## Abstract

Gene expression analyses to study the development of cotton fibers require high-quality nucleic acid. The conventional methods of nucleic acid extraction results in sub-quality nucleic acids with low yields. Young fibers are rich in polyphenols and sugars that react with nucleic acid to form phenols and insoluble substances. Furthermore, mature fibers contain more than 95% cellulose, hindering the nucleic acid isolation. Cytoplasm collapse and cellulose deposition also result in a very low yield of nucleic acid. Three different methods of RNA isolation from different cotton tissues were compared in this study to determine the best and most efficient one. The integrity and quality of RNA were analyzed using UV spectrum, agarose gel electrophoresis, RIN values, PCR, and Northern blot hybridization. RNA of functional quality was observed when using the high ion and pH method, with an A260/A280 ratio up to 1.87 and an average yield of 0.68 mg g^-1^ from fiber cells. From leaves, we found an A260/A280 ratio of 2.02 and an average yield of 6.35 mg g^-1^, which is suitable for molecular biology experiments. The extraction buffer with a high ion density and pH value include Tris-HCl, LiCl, EDTA, SDS, sodium deoxycholate, Nonident P-40, mercaptoethanol, and PVP. The addition of sodium deoxycholate and Nonider-40 (NP-40) enhances the density of other salt compounds and elevates the pH value. The results depicted that the high ion and pH method is a simple and effective way to extract a copious amount of high-quality RNA from polysaccharide-rich tissues. This method is also suitable for the extraction of cotton genomic DNA with high purity. Genomic DNA extracted from cotton using this method showed an A260/A230 ratio up to 2.09 and a yield of 1.44 mg g^-1^. This method is useful for isolating DNA and RNA from cotton fibers and produces high yields and quality at a comparatively low cost.

## INTRODUCTION

Cotton is the world’s most economically important cash crop to produce natural textile fibers. To improve the yield and quality of cotton it is necessary to research the molecular mechanisms of cotton growth and development. For studies it require the isolation of high-quality RNA and DNA [1,2]. In cotton fibers, the cellulose contents are particularly high and are the main component of cell wall synthesis. During cell development, the vacuole gradually increases and occupies most of the cell volume, which contains large amounts of secondary metabolites, such as polyphenols, terpenes, and tannins [3,4]. During nucleic acid (RNA) extraction from fibers, the breakdown of large vacuoles releases acidic materials like polyphenols, lipids, proteins, nucleic acids, and insoluble polysaccharides, which can cause degradation of the nucleic acids by various irreversible reactions [5–7]. Extraction of proteins and nucleic acid from young as well as mature fibers is a very tedious job because young fibers are rich in polyphenolics and polysaccharides and mature fibers contain little cytoplasm. Commercial kits are available for nucleic acid extraction from various plant tissues, but no effective kit is available for RNA extraction from young and mature cotton fibers. DNA of sufficient quality can be harvested from plant tissues using the existing kits, but the yield is often low [2,8].

Different kinds of naturally occurring RNAs in plant tissues include large amounts of rRNA (80-90%) and tRNA but very low levels of mRNA (5-10%). Due to the small amount of mRNA, it is very difficult to isolate it in a functional form from the mature cotton fiber. Conventional methods of extraction result in reddish-brown, insoluble RNA that is difficult to use for enzymatic reactions and hence is not suitable for fiber gene regulation and expression studies [9–11]. Although many extraction methods are used to extract RNA from cotton materials, the purity and yield of RNA extracted from cotton fibers are inadequate due to the reasons discussed above. During DNA extraction from cotton, phenolics, and sugars interfere, which hinders the extraction of clean genomic DNA. Currently, CTAB based nucleic acid extraction method is frequently used by researchers. This method can exclude sugars and phenol materials in addition to improving the quality of DNA due to the presence of antioxidant components such as PVP [12,13].

Different methods of RNA extraction have broad applications depending upon tissue composition. The TRIzol^®^ Reagent method is simple, fast, and has efficient RNase inhibitory activity, and this method can be used for RNA extraction from a wide variety of plants. The guanidinium isothiocyanate method is the classic method for RNA extraction, with some modifications according to different cell types. However, repeated experiments have proved that these two methods are unsuitable for RNA extraction from cotton fibre cells. The current effort was provoked by the need for an improved method to harvest an ample amount of high-quality RNA from polysaccharides and high gossypol matrices that can be used in gene expression studies. We used a high ionic strength and pH buffer along with antioxidant PVP, Nonident P-40, and mercaptoethanol to improve the RNA extraction efficiency and preparation of high-yield and high-purity RNA. The antioxidant inhibited the oxidation of phenols and selective precipitation under high salt conditions, which eliminated the interference of sugars with RNA, with minimum chances of cross-contamination. Further work needs to be carried out using this protocol to explore methods of extracting cotton genomic DNA and cloning gene fragments for downstream processes. Regarding per sample cost of extraction of RNA through kits was found to be ranged between $6.00 to $10.0 by using different kits e.g., Qiagen RNeasy mini kit as compared to $1.00/sample by using the High Ion and pH method.

## MATERIALS AND METHODS

### Plant materials

Four cotton lines were used in the experiment, the first was a commercial cotton variety Xuzhou 142 (upland cotton) with normal fibers, and the second and third were two fiber mutants of Xuzhou 142 (Xuzhou 142-N without fuzz that is also called naked seed or fuzz-less mutant, and Xuzhou 142-fl without lint and fuzz (that is also called as a fibreless mutant), and the fourth was Ligon lint-less mutant with very short fibers (only about 6~8mm length). Etiolated seedlings and bolls of cotton and leaves of tobacco variety “Hongda” were prepared for RNA and DNA isolation. RNA was isolated from a cotton embryo, seed coat, and fiber at 15 DPA. Samples were collected in ice and immediately frozen in liquid nitrogen and stored at −70°C.

### RNA extraction

#### RNA extraction by the high ionic and pH method

All solutions used for RNA extraction were sterilized at high pressure and treated with diethylpyrocarbonate (DEPC). Glassware was sterilized at 200°C for 6 h and soaked in 0.1% DEPC for 30min to inhibit RNase activity. Extraction buffer was prepared by adding Tris-HCl (200mmol.L^-1^) pH 8.5, LiCl (300mmol.L^-1^), EDTA (10mmol.L^-1^), 1.5% SDS, 1.5% sodium deoxycholate, 1.5% Nonident P-40, treated with DEPC and stored at −20°C. Mercaptoethanol (75mmol.L^-1^) and 1% PVP (w/v) were added just before use.

Grind 0.35g of plant materials to the fine powder with liquid nitrogen in a pre-cooled pestle and mortar. Transfer the powder to 15ml centrifuge tubes, add 5ml of extraction buffer, and mix by inverting the tube. Add 5ml chloroform and mix, placed at room temperature for 5-10min for denaturation of proteins, and then centrifuge at 9500xg for 15min at 4°C. Add chloroform again if protein exists in the material. Transfer the supernatant to a new 15ml centrifuge tube and add 1/10 volume 3 mol.L^-1^ NaCl and two times the volume of ethanol then precipitate at −20°C for 1h. Centrifuge at 4000xg for 15min at 4°C. Add 500μl TE and centrifuge at high speed for precipitation. Add 1/10 volume 3mol.L^-1^ NaCl and 275μl isopropanol and centrifuge at 12000xg for 15min at 4°C. Precipitate by adding 70% ethanol and dry at room temperature. Dissolved the precipitate in 300μl of TE and add 100μl of 8mol.L^-1^ LiCl, placed in the ice bath overnight. Then centrifuge at 4°C, 12000xg for 10 min, dissolve the pellet in 300μl of TE and add 30μl 3mol.L^-1^ NaCl and 660μl of ethanol, precipitate at −20°C for 1h. Centrifuge at 4°C, 12000xg for 10min. Washed the pellet completely with 70% ethanol, and dried at room temperature. The pellet was dissolved in an appropriate volume of TE and stored at −20°C.

#### RNA extraction by the Trizol method

A total of 0.35g of plant tissue was pulverized, and RNA was extracted by using Trizol Reagent (Takara, Shanghai, China) according to the manufacturer’s instructions provided with the kit.

#### RNA extraction by guanidinium isothiocyanate method

RNA was extracted by the guanidinium isothiocyanate method using 0.35g plant tissue as described by Wadsworth *et al.* [14].

### DNA Extraction by a high ionic and pH method

Grind 0.35g of plant tissue materials to the fine powder with liquid nitrogen in a pestle and mortar. Transfer the powder to 15ml centrifuge tubes, add 5ml of extraction buffer, and mix by inverting the tubes. Add 5ml chloroform and mix, placed at room temperature for 5-10min for denaturation of proteins, centrifuge at 9500xg for 15min at 4°C. Add chloroform again if protein exists in the material. Transfer the supernatant to a new 15ml centrifuge tube and add 1/10 volume 3mol.L^-1^ NaCl, and twice times the volume of ethanol and precipitate at −20°C for 1h. Centrifuge at 4000xg for 15min at 4°C. Add 500μl TE and centrifuge at high speed for precipitation. Add 1/10 volume 3mol.L^-1^ NaCl and 275μl isopropanol and centrifuge at 12000xg for 15min at 4°C. Precipitate by adding 70% ethanol and dry at room temperature. Dissolved the pellet in 500μl TE, add an equal volume of phenol: chloroform: isoamyl alcohol, and mix by inverting the tube gently. Centrifuge at 12000xg for 10min at 4°C, transfer the supernatant to a new centrifuge tube, repeat this step, and add chloroform and mix. Add 30μl of 3mol.L^-1^ NaCl and 660μl ethanol, precipitate at −20°C for 1h. Centrifuge at 12000xg for 10min at 4°C. Wash the pellet with 70% ethanol, and dry at room temperature. Dissolve in an appropriate volume of TE saved at −20°C.

### Quality analysis of RNA and DNA

RNA and DNA were quantified by using Nanodrop 2000 (Thermo Fisher Scientific, USA) as reported previously [15]. The quality was also determined through agarose gel electrophoresis. RNA Integrity Number (RIN) was determined through the Agilent 2100 Bioanalyzer system using RNA 6000 Pico kit (cat # G2939BA) following the manufacturer’s instructions.

#### cDNA synthesis and PCR

Clean RNA is a prerequisite for the construction of full-length cDNA as reverse transcription is very sensitive to impurities. So, RNA quality was verified by reverse transcription and PCR amplification. The first strands of cDNA were synthesized with 1μg RNA as a template according to the Takara cDNA synthesis kit (Cat#6130). Synthesis of cDNA was accompanied at 42°C for 30min followed by 70°C for 10min. Sucrose synthase (SuS) and Expansin gene were amplified from cDNA using gene-specific primers (*SuS* 5’-CTTGAGAAATTCTTGGGTACTATCC-3’; 5’-CTCGGTGTAAGGGAAATAAATAGAC-3’: *EXP* 5’-AGTCGAACCATAACCGTGAC AGCC-3’; 5’-CCCAATTTCTGGACATAGGTAGCC-3’).

PCR reaction mixture was prepared as: 2μl PCR buffer (10x), 2μl dNTPs (1mM), 2.5 units Taq polymerase, 2μl Primer 1 (10mM), 2μl Primer 2 (10mM), 1.5μl DNA (about 25ng) template. The final volume was adjusted up to 20μl by adding water. PCR reaction was performed by using the following PCR conditions: 95 *°*C initial denaturation for 4min, 95 °C 30s, 55 °C 45s, 72°C 45s, 35cycles, 72°C 7min. The amplified product was separated on 1.0% agarose gel and visualized by ethidium bromide staining.

#### Northern Blot Hybridization

Total RNA isolated was denatured and separated by agarose gel electrophoresis. Northern blotting was performed as previously described by Ruperti et al. [16] and blots were hybridized with a radioactively labeled expansin gene with α-^32^P as a probe. A standard protocol was used for hybridization and washing [17].

#### Restriction Endonuclease Cleavage of DNA extracted by high ion and pH method

A total of 10μg of DNA was digested with restriction enzyme (1U/μg DNA) in the recommended buffer at 37°C for 1hr. The digested DNA and α-^32^P labeled DNA marker were electrophoresed on a gel and transferred to a nylon membrane.

#### PCR Amplification from DNA

PCR was performed by using genomic DNA as a template. Set of primers used to amplify *CeLA* (cellulose synthase), *EXP* (expansion), *EDOGT* (wood-glucan o-methyltransferase) and *E14G* (ß-1,4-ß-glucanase), *PEPCase* (Phosphoenol Pyruvate Carboxylase) genes are given in table 1. PCR reaction mixture was prepared as: 2.5μl (10xPCR buffer), 0.5μl (10mM dNTPs), 1.25 units Taq polymerase, 1μl (10mM Primer 1), 1μl (10mM Primer 2), 1μl DNA (about 25ng) template, total volume 25μl was adjusted by adding water. PCR reaction conditions: 95°C initial denaturation 5mins, 95°C 45s, 56°C 45s,72°C 45s, 35cycles,72°C 10min.

**Table 1:**
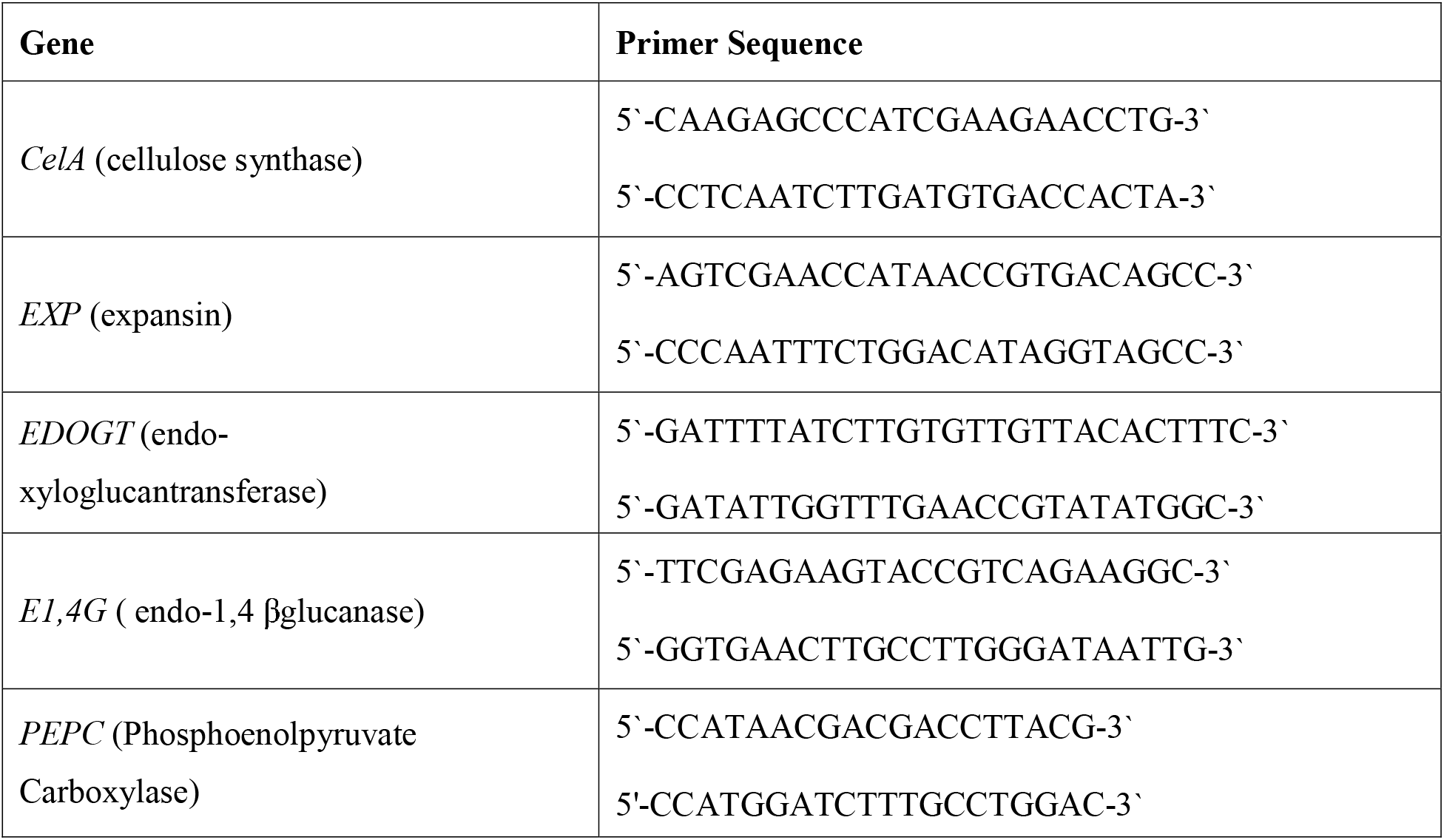
Primers sets used for gene amplification

## RESULTS

### Purity and Yield of RNA and DNA

#### Purity and yield of RNA extracted by three methods

We compared RNA purity and yield between the three extraction methods. The TRIzol^®^ Reagent method is the best for RNA extraction from tobacco leaves as its purity and yield are highest when using this protocol out of the three methods that we tested, but it is not suitable for RNA extraction from cotton tissues. The A260/A280 ratio is significantly lower than in the high ion and pH buffer method. Similarly, the guanidinium isothiocyanate buffer extraction method is not suitable for small amounts of RNA extraction from cotton ovule tissues as the A260/A280 < 1.52 and the yield is relatively low due to protein contamination. The high ion and pH extraction buffer method is best for RNA extraction from cotton ovules as purity was up to 2.0 and the yield was higher than that of the other two methods (Supplementary Table S1).

### Analysis of RNA and DNA Quality

#### Gel electrophoresis and analysis of RNA purity

We compared different RNA extraction methods using gel electrophoresis. The appearance of two sharp bands of 28S and 18S RNA on agarose gel electrophoresis with ethidium bromide (EB) staining indicates the intactness of RNA. Using the high ion and pH buffer solution with PVP and ß-mercaptoethanol, RNA isolated from cotton leaves and fibers showed two distinct and intact 28S and 18S sharp bands which indicate the presence of ribosomal RNA subunits (Fig. 1). RNA has selective precipitation in a LiCl solution, so there are no 5S RNA molecules. Using this method of extraction without PVP and β-mercaptoethanol, degradation of cotton leaf RNA occurs, but remnants are smaller in the loading well. PVP and β-mercaptoethanol can inhibit the oxidation of polyphenols and prevent their co-precipitation with nucleic acids as well as irreversible denaturation of RNases. A high ion and pH buffer solution helps to eliminate the irreversible binding of intracellular polysaccharides and polyphenols with nucleic acid. The RNA extracted by the TRIzol^®^ Reagent method was electrophoresed on the gel with two bands, but a copious number of proteins and other impurities remain in the loading well. RNA extracted by guanidinium isothiocyanate buffer from cotton fibers remains in the loading well and did not show the two sharp bands. The TRIzol^®^ Reagent method and guanidine isothiocyanate extraction cannot eliminate interfering phenolics and polysaccharides from the cotton fiber cells which leads to degradation and RNA becoming insoluble in TE solution. The electropherogram generated by using Agilent 2100 Bioanalyzer revealed intactness and stability of RNA isolated by high ion and pH method with RIN values above 9 (Fig. 2).

**Fig. 1.**
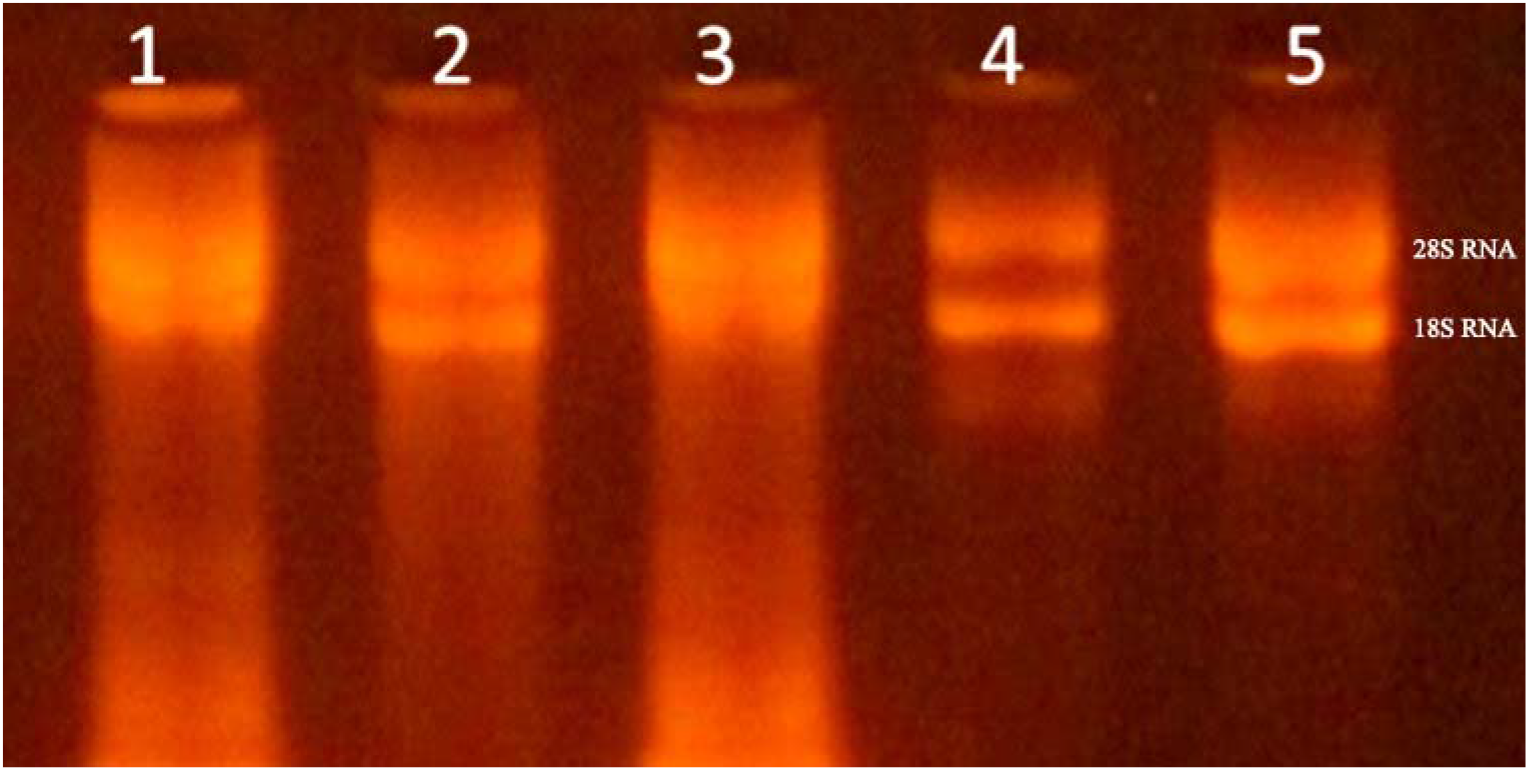
Agarose gel electrophoresis of RNA extracted by three methods. 1: RNA of cotton fibre extracted by guanidine isothiocyanate method, 2: RNA of cotton fibre extracted by Trizol methods, 3: RNA of the cotton leaf by the high ion and pH method without adding PVP and β-mercaptoethanol, 4: RNA of cotton leaves by the high ion and pH method. 5: RNA of cotton fibre extracted by the high ion and pH method.

**Fig. 2.**
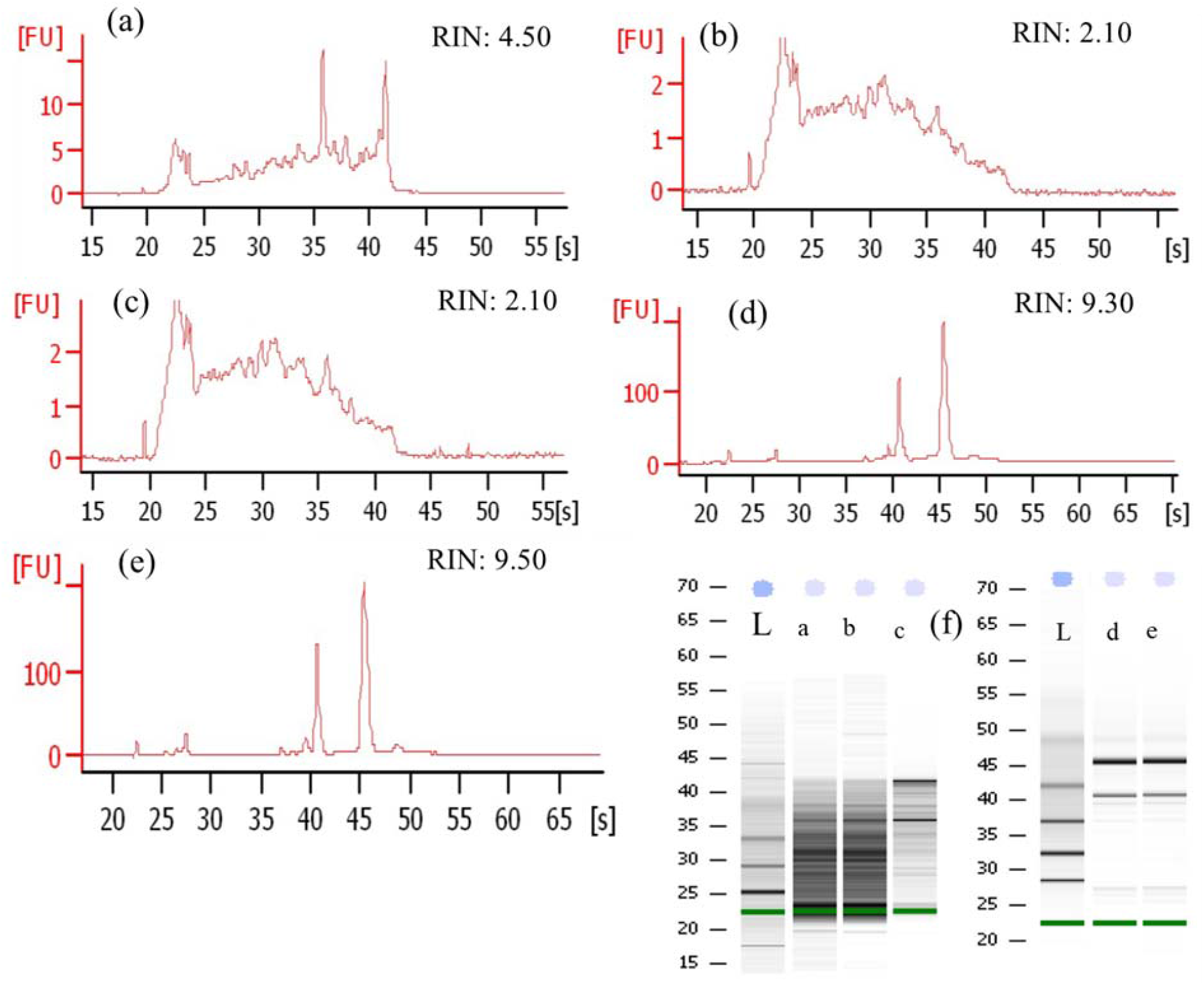
Electropherogram of RNA samples showing RIN values. Samples range from degraded (RIN<5) to intact (RIN≥6); (a) RNA of cotton fibre extracted by guanidine isothiocyanate method; (b) RNA of cotton fibre extracted by Trizol methods; (c) RNA of the cotton leaf by the high ion and pH method without adding PVP and β-mercaptoethanol; (d) RNA of cotton leaves by the high ion and pH method; (e) RNA of cotton fibre extracted by the high ion and pH method; (f) RNA samples (a-e) analyzed on the Agilent 2100 Bioanalyzer system through capillary gel electrophoresis L: RNA ladder 15S-70S.

For the UV spectral analyses, we first compared the UV spectra of RNA that were extracted by different extraction methods. UV spectral analysis of tobacco RNA extracted by the TRIzol^®^ Reagent method has full absorption at 200–300nm. At 260nm it showed a maximum absorption peak, whereas at 230nm and 280nm the absorption value was significantly lower than at 260nm. This indicates that the RNA extracted was less polluted with proteins and other salts. However, the absorption spectra of RNA that was extracted from cotton tissues showed an unusual trend where its absorption peak transferred to 270nm, which indicated contamination of phenolic materials. At 230nm for cotton leaf RNA that was extracted by guanidinium thiocyanate liquid extraction, absorption values are higher than the high ion and high pH method, which shows that these methods of RNA extraction are not suitable for cotton due to the contamination of salts such as guanidinium thiocyanate and phenols. The absorption spectra showed that guanidinium isothiocyanate and other salt materials polluted cotton fiber RNA that was extracted using guanidinium isothiocyanate. Therefore, this method is not suitable for RNA extraction from cotton fibers. RNA from cotton fibers that were extracted with the high ion and pH buffer solution showed a complete absorption, which indicates that RNA that was isolated by this method is less polluted by salt substances such as proteins and polysaccharides and is suitable for cotton fiber RNA extraction.

### RT-PCR and Northern Hybridization

We performed qRT-PCR and Northern hybridization reaction tests to validate the purity and integrity of cotton RNA that was isolated using the high ion and pH buffer method. The results showed that RNA from the high ion and pH buffer solution meets the requirements of enzymatic reactions, and thus is suitable for general molecular biology experiments. As shown in Fig. 1, RNA extracted from different materials can be used for cDNA synthesis. Using cDNA as a template SuS and Expansion genes were amplified, and a sharp amplicon was seen from RNA isolated from the high ion and pH buffer method. PCR results demonstrated that there are significant differences in the performance of each method in terms of PCR amplification. The RNA isolated by other extraction methods failed to produce detectable amplification products (Fig. 3). These genes play an essential role in all stages of plant development. Using the expansin gene with α-^32^P as a probe distinct and sharp hybridization signal without smear was seen (Fig. 3c). The results indicate the good integrity of RNA for downstream molecular procedures.

**Fig. 3.**
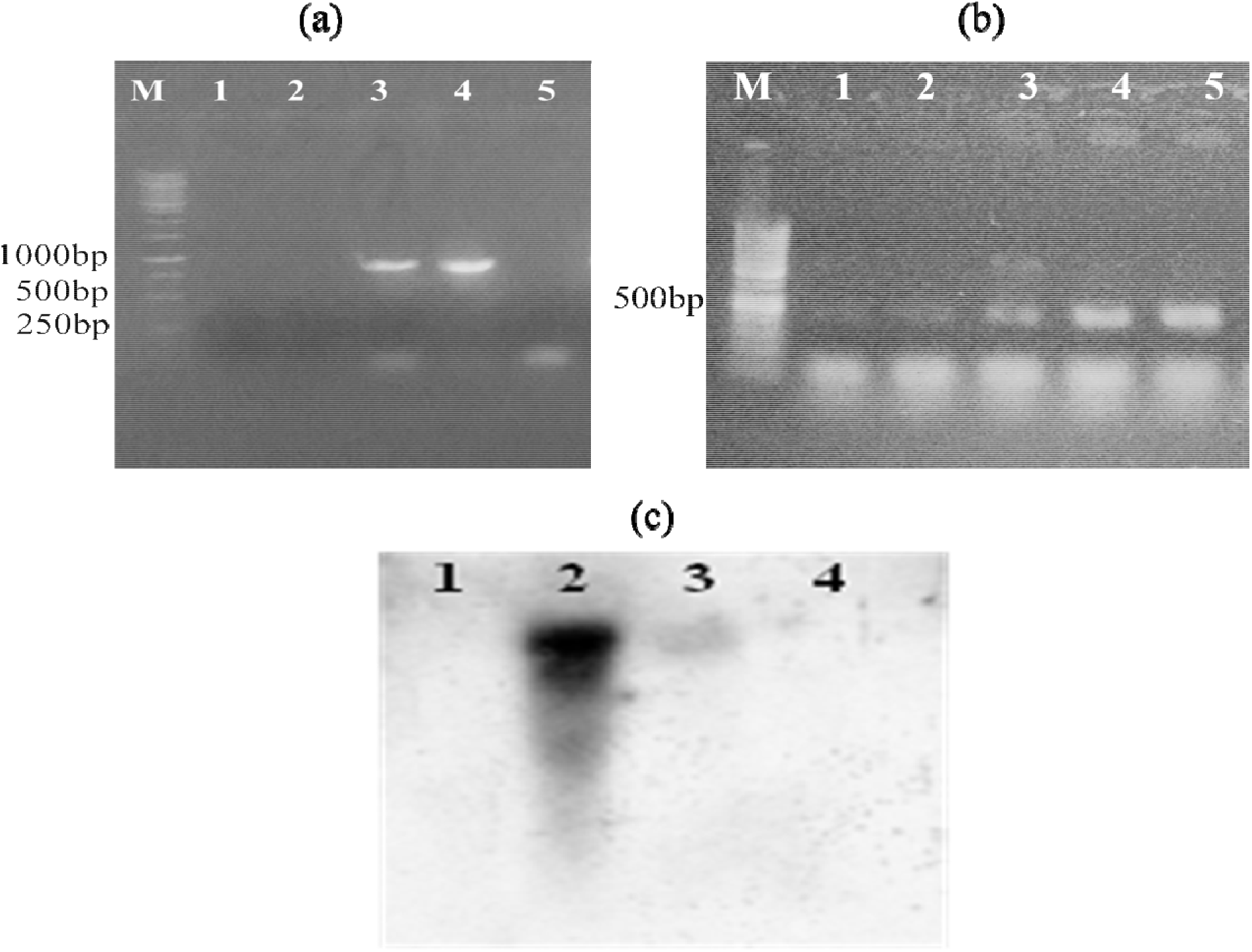
Electrophoretic analysis of PCR products amplified from first strand cDNA and purified total RNA isolated by different methods as template; (a) PCR amplification of SuS; M: DNA marker Lane 1: PCR using RNA isolated from cotton fiber by guanidine isothiocyanate method, 2: PCR using RNA isolated from fiber by Trizol methods, 3, 4: PCR using RNA isolated from leaf and fiber respectively by the high ion and pH method, 5: PCR using RNA isolated from leaf by the high ion and pH method without adding PVP and β-mercaptoethanol; (b) PCR amplification of Expansin gene M: DNA marker Lane 1: PCR product amplified from cDNA made from RNA isolated from cotton fiber by guanidine isothiocyanate method, 2: PCR product using RNA isolated from fiber by Trizol methods, 3: PCR using RNA isolated from leaf by the high ion and pH method without adding PVP and β-mercaptoethanol, 4, 5: PCR using RNA isolated from leaf and fiber respectively by the high ion and pH method; (c) Northern blot hybridization using RNA isolated through high ion and pH method; 1, 2, 3 and 4: RNAs from 5, 10, 15 and 20DPA fiber cells respectively hybridized with a probe (expansin gene).

### Gel electrophoresis and analysis of DNA purity

We performed agarose gel electrophoresis of genomic DNA isolated from different cotton materials by the high ionic and pH buffer method. The results showed sharp and clear bands with less contamination when fractionated on a 1.0 % agarose gel (Fig. 4a). UV spectral analysis revealed a complete absorption spectrum with a maximum absorption peak occurring at 260nm along with 230nm values significantly lower than those at 260nm and 280nm. These results showed less protein and salt contamination. The value of A260/230 increased above 1.9 with an average yield above 1mg.g^-1^ from all three used cotton materials (Supplementary Table S2).

**Fig. 4.**
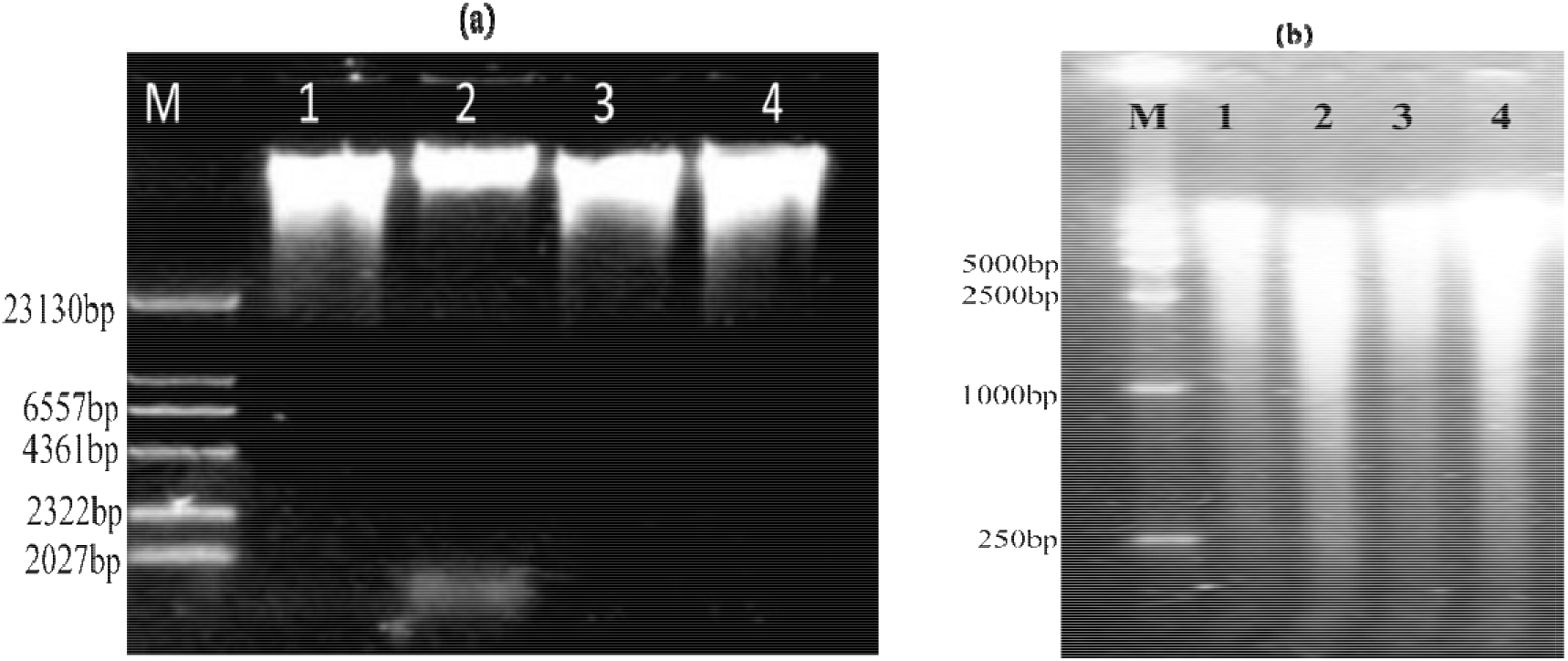
(a) Agarose gel electrophoresis of genomic DNA; (b) Agarose gel electrophoresis of genomic DNA digested with *EcoRI*. M - DNA marker. 1, 2, 3 and 4: DNAs from 142 Xuzhou standard type, Xuzhou 142-N mutant, Xuzhou 142-fl mutant, and Ligon-lintless mutant, respectively.

#### Restriction enzyme cleavage of DNA and Southern Hybridization

After digestion with restriction enzymes, gel electrophoresis showed a uniform dispersion with a small amount of DNA remaining in the loading well (Fig. 4b). Different fragments of *EDOGT* and *EXP* genes were detected through Southern hybridization. The appearance of different-sized fragments speculate the presence of *EDOGT* and *EXP* gene families in the cotton genome. These results indicate that the purity and integrity of DNA are consistent with no shearing and contamination (Fig. 5).

**Fig. 5.**
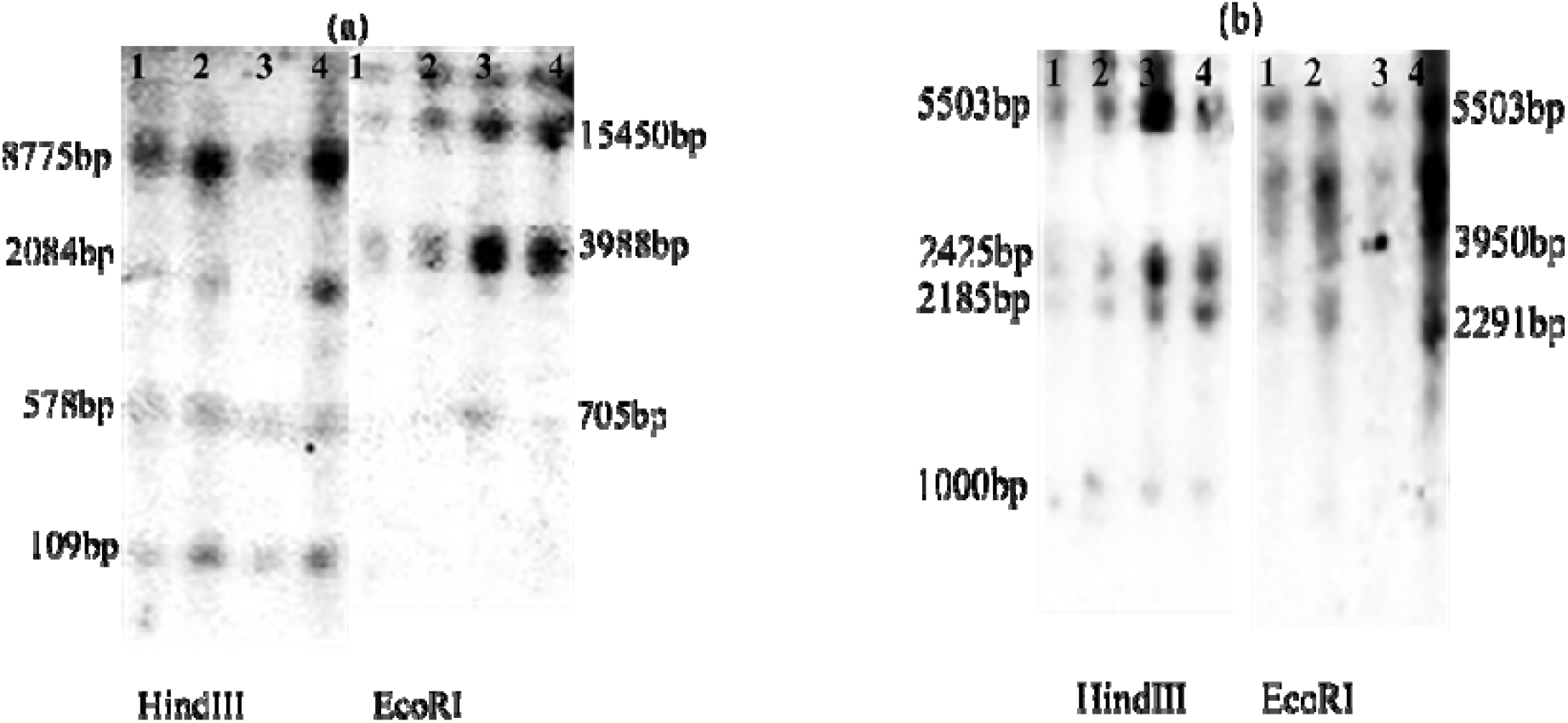
(a) Southern hybridization of *EDOGT* (Endoxyloglucan transferase); (b) Southern hybridization of Expansin gene; 1: Xuzhou 142 standard, 2: Xuzhou 142-N mutant, 3: Xuzhou 142-fl mutant, 4: Ligon-lintless cotton. Approximately 10 μg genomic DNA of each sample was digested with HindIII and EcoRI, and the Endoxyloglucan transferase gene and Expansin gene fragment were used as a probe.

### PCR amplification from DNA

The PCR was performed using genomic DNA that was isolated from yellow seedlings of Xuzhou 142 normal cotton as a template. Five genes *CeLA* (cellulose synthase genes), *EXP* (Expansin), *EDOGT* (Endoglucanase transferase) and *E1, 4G* (Endo1, 4-β-glucanase), *PEPC* (Phosphoenol pyruvate Carboxylase), respectively were successfully amplified using DNA extracted from the high ion and pH buffer method (Fig. 6). Therefore, the results of genomic DNA meet the requirements of the PCR reaction.

**Fig. 6.**
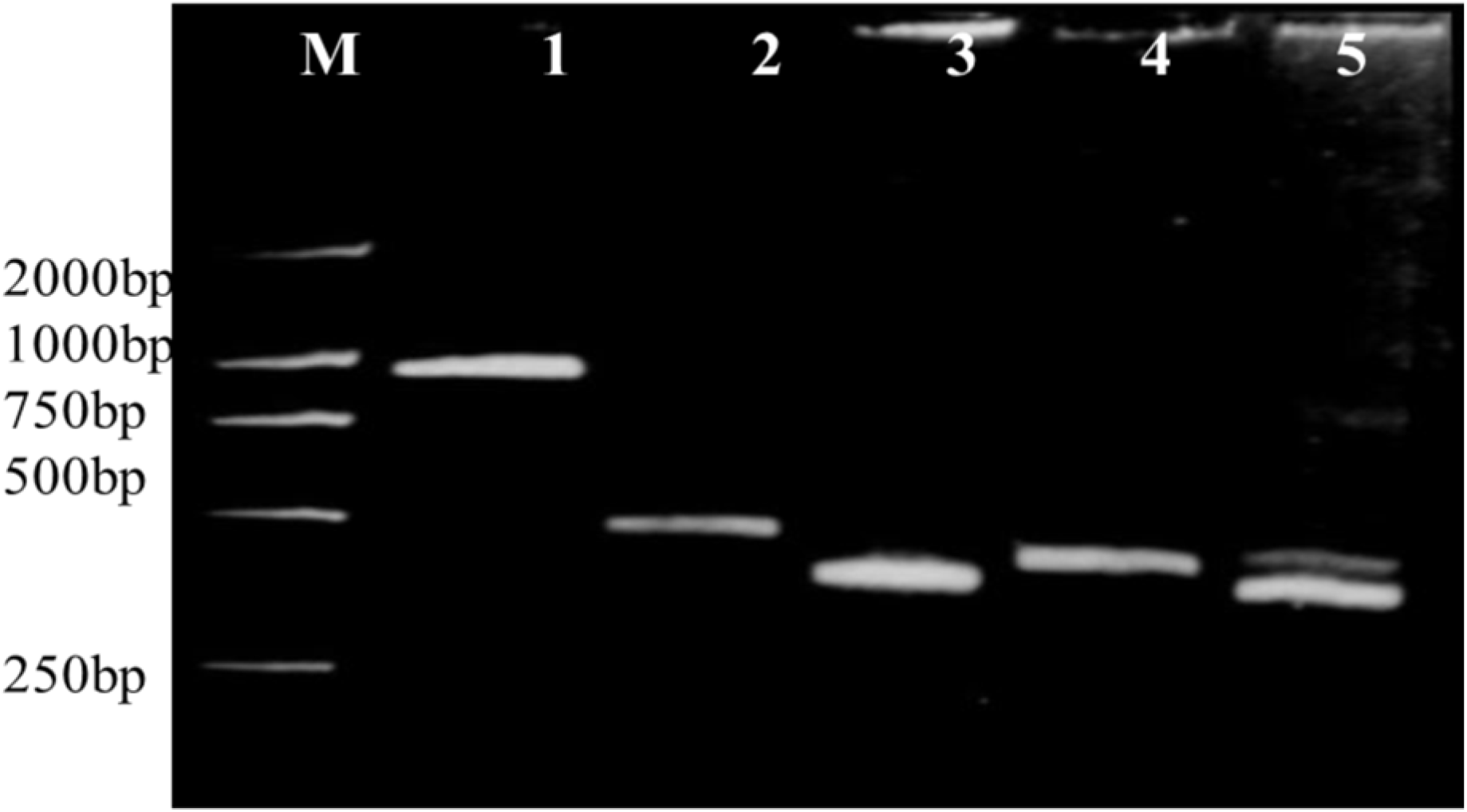
PCR amplification of different genes from genomic DNA isolated using high ion and pH method; M: DNA marker; 1, 2, 3, 4 and 5: PCR products of *CelA, EDOGT, EXP, PEPC* and *E14G* gene, respectively.

## DISCUSSION

This study has established a simple and feasible method of RNA extraction from cotton fibers, embryos, seed coats, and leaves after repeated experiments. Previously, RNA was isolated from different cotton tissues using different methods, but degradation and low yield and purity generally affected the samples (Table 2). The extraction method described in this paper is suitable for extracting RNA from various cells of the cotton ovule. The success of this protocol is attributable to the following key points: 1) the ratio of extraction buffer to plant material and the ionic strength and pH of the buffer was improved, and the addition of detergents, PVP, and mercaptoethanol can fully prevent phenolics and sugar materials from combining with nucleic acid, while EDTA and SDS inhibit endogenous RNase activity. SDS completely solubilizes and denatures the proteins. RNA degradation occurred when using an extraction buffer without PVP and mercaptoethanol (Fig. 1). 2) The combined use of phenol or phenol and chloroform can prevent the partitioning of RNA into the organic phase and nucleic acid, and hence improves the yield of RNA compared to using chloroform alone for RNase denaturation and to remove homogenate proteins during centrifugation. 3) Salts help to remove polysaccharides, so we used salts for selective precipitation with ethanol and we used high-speed centrifugation to remove the insoluble part; therefore, the impact of polysaccharides on nucleic acid can be reduced to avoid DNA contamination. PVP and EDTA both function to inhibit the oxidation of polyphenols in cell walls and the extracellular matrix. EDTA also has a role in the inactivation of DNAse by binding to metal cations that are essential for enzyme activity [18,19]. β-mercaptoethanol acts as a strong reducing agent that prevents any oxidation reactions and irreversibly denatures RNases [20]. Phenol-chloroform makes phase separation easier (due to the difference in density) as centrifugation results in the formation of two phases, the lower organic phase, and the upper aqueous phase. The combination of phenol-chloroform is more effective in denaturing proteins and reduces the formation of an insoluble RNA-protein complex at the interphase. Phenol keeps about 10-15% of the aqueous phase, which results in the loss of some RNA, while chloroform stops this retention of water and therefore improves RNA yield [21].

**Table 2:** Comparison of previous studies of RNA extraction from cotton tissues

| Method Used | Cotton Tissue | A260/280 | Reference |
| --- | --- | --- | --- |
| Lithium Dodecylsulphate | Leaf, pollen | N A | [26] |
| Guanidine Isothiocyanate | Stem, root, fiber, leaf, ovule, flower | N A | [3] |
| Hot borate | Ovules, petal, anthers, leaf, embryo, root | 1.79-1.80 | [12] |
| Improved phenol sodium dodecyl sulfate | Leaves, flowers, and buds | 2.0-2.2 | [27] |
| CTAB acidic phenol | Leaves, roots, anthers, hypocotyls, and fibers | 1.90-2.0 | [28] |
| CTAB-PVP-LiCl | Fibers, leaves, Petals, radicals, hypocotyls, ovule and anther | 1.74-1.98 | [29] |
| Mini scale hot borate method | Leaves, anthers, -1, 0, 5, 8 and 10 DPA fibers | 1.77 - 2.13 (at 260/230) | [30] |
| CTAB Ammonium acetate | Leaves, stem, ovule, petals, and fiber | 1.80-1.85 | [1] |
| Used existing methods with modifications | Cotton roots | 1.90 | [5] |
| The High ion and pH buffer method | Cotton fiber, leaf, embryo, seed coat | 1.87-2.09 | Current study |

N A: Not Applicable

The TRIzol^®^ Reagent method of RNA extraction is very effective for tobacco and other types of plants with less sugar, but it is not suitable for cotton as RNA appears brown and degradation occurs. The brownish color and degradation of RNA may be due to polysaccharides, proteins, and the oxidation of phenolics that irreversibly interact with RNA [6]. The TRIzol^®^ Reagent cannot effectively eliminate phenols, polysaccharides, and other salts that are abundant in fibers. The oxidation of polysaccharides and other secondary metabolites co-precipitate with nucleic acids upon cell lysis and forms an opaque slurry that leads to variations in ultraviolet absorption spectrum results [22]. These anomalies in the UV spectrum may be caused by the degradation of nucleic acids. Similarly, the guanidinium isothiocyanate extraction method is not suitable for RNA extraction from cotton ovules and is hazardous due to toxic substances that are present in it. It is useful for RNA extraction from cotton leaves but not practically important for cotton fibers due to the presence of endogenous polysaccharides and sugars that have been found to co-purify during RNA precipitation. Therefore, RNA purity and yield are less compared to the high ion pH extraction buffer protocol. Thus, the guanidinium isothiocyanate and TRIzol^®^ Reagent methods are not appropriate for RNA extraction from the cotton ovule as this tissue is rich in phenols and sugars. He et al. [23] isolated RNA from Walnut buds using four methods: CTAB, Trizol, D326A reagent, and the RP3301 kit. There was a certain degree of RNA degradation by the CTAB, TRIzol^®^, and D326A reagents along with proteins, polyphenols, and polysaccharide impurities. The results of the current study also coincide with previous findings that acid guanidinium thiocyanate, SDS-phenol, and/or a hot borate method remained unsuccessful for RNA isolation from the tissues that are rich in secondary metabolites [24,25]. Cotton genomic DNA that was extracted using the high ion and pH buffer method proved to be of high quality and meets the requirements of molecular experiments. Genomic DNA extraction with this method from the four cotton materials showed the same results in DNA purity, yield, and DNA digestion with restriction enzymes.

The high ion and pH method described in this study allowed the extraction of intact, high-yield, and high-quality RNA from leaves and fiber samples of cotton plants. It is obvious from the results that this method can be used for other plant tissues that have high levels of phenolics and secondary metabolites from which RNA isolation is notoriously difficult. The harvested RNA can be used for qRT-PCR and other downstream molecular biology experiments. This method is efficient and simple and does not require specialized equipment. The high ion and pH method will pave the way toward the development of commercial kits for the extraction of high-quality RNA from cotton fibers.

## Abbreviations

CTAB: Cetyl Trimethyl Ammonium Bromide
DNA: Deoxyribose Nucleic Acid
DPA: Days Post Anthesis
EDTA: Ethylene Diamine Tetra Acetate
mg g-^1^: milligram per gram
PCR: Polymerase Chain Reaction
PVP: Polyvinylpyrrolidone
RIN: RNA Integrity Number
RNA: Ribose Nucleic acid
RT-PCR: Real-Time PCR
SDS: Sodium Dodecyl Sulfate
TE: Tris EDTA
uL: microliter
UV: UltraViolet.

## ACKNOWLEDGMENTS

This work was supported by the National Natural Science Foundation of China and Zhejiang Province Science Research Project (31671763, 31471567, and 2017C32077) and the Higher Education Commission (HEC), Pakistan.

## LIST OF TABLES

**Supplementary Table S1:** Comparison of yield and purity of RNA from different plant tissues by three methods

| Method | Plant Tissue used | Purity $A_{260}/A_{280}$ | Average | Yield /mg.g<sup>-1</sup> | Average yield /mg.g<sup>-1</sup> |
| --- | --- | --- | --- | --- | --- |
| The High Ion and pH method | Tobacco leaf | 1.48~1.89 | 1.653±0.2<br>12 | 5.32~5.80 | 5.59±0.24 |
|  | Cotton leaf | 1.92~2.02 | 1.973±0.5 | 6.04~6.66 | 6.35±0.31 |
|  | 15DPA cotton fiber | 1.58~1.87 | 1.723±0.1<br>45 | 0.42~0.97 | 0.68±0.27 |
|  | 15DPA cotton embryo | 1.70~1.98 | 1.843±0.1<br>40 | 1.07~1.27 | 1.09±0.11 |
|  | 15DPA<br>cotton seed<br>coat | 1.70~2.00 | 1.863±0.1<br>52 | 0.61~0.90 | 0.70±0.14 |
| TRIZOL<br>method | Tobacco<br>leaf | 1.89~1.95 | 1.927±0.0<br>32 | 5.89~7.61 | 6.75±0.86 |
|  | Cotton leaf | 1.36~1.47 | 1.413±0.0<br>55 | 1.89~2.37 | 2.13±0.24 |
|  | 15DPA<br>cotton<br>fiber | 1.33~1.42 | 1.380±0.0<br>46 | 0.34~0.52 | 0.42±0.09 |
|  | 15DPA<br>cotton<br>embryo | 1.38~1.45 | 1.410±0.0<br>36 | 0.55~0.72 | 0.65±0.08 |
|  | 15DPA<br>cotton seed<br>coat | 1.24~1.52 | 1.403±0.1<br>46 | 0.43~0.56 | 0.52±0.06 |
| Guanidine<br>Isothiocyana<br>te<br>method | Tobacco<br>leaf | 1.89~1.95 | 1.913±0.0<br>32 | 4.40~5.21 | 4.81±0.40 |
|  | Cotton leaf | 1.85~2.01 | 1.933±0.0<br>80 | 3.21~3.72 | 3.47±0.25 |
|  | 15DPA<br>cotton<br>fiber | 1.21~1.32 | 1.267±0.0<br>55 | 0.23~0.31 | 0.27±0.04 |
|  | 15DPA<br>cotton<br>embryo | 1.35~1.52 | 1.443±0.0<br>86 | 0.45~0.62 | 0.53±0.08 |
|  | 15DPA<br>cotton seed<br>coat | 1.35~1.46 | 1.407±0.0<br>55 | 0.48~0.55 | 0.51±0.03 |

The results are an average of three times RNA extraction and standard deviation, DPA means days post anthesis.

**Supplementary Table S2:** Comparison of purity and yield of DNA from different cotton seedlings by high ion-density and pH method

| Method | Varie<br>ties<sup>(1)</sup><br>(seed<br>ling) | Purity<sup>(2)</sup><br>$A_{260/230}$ | Average | Yield<sup>(2)</sup><br>/mg·g<sup>-1</sup> | Average<sup>(2)</sup><br>yield<br>/mg·g<sup>-1</sup> |
| --- | --- | --- | --- | --- | --- |
| The High<br>ion and<br>pH<br>method | Xuzh<br>ou<br>142<br>(nor<br>mal<br>type) | 1.96~2.09 | 2.027±0.065 | 1.32~<br>1.5 | 1.44±0.09 |
|  | Xuzh<br>ou<br>142-N | 1.99~2.06 | 2.027±0.035 | 1.1~1.<br>58 | 1.33±0.24 |
|  | Xuzh<br>ou<br>142-fl | 1.94~2.65 | 2.330±0.36 | 1.21~<br>1.55 | 1.35±0.17 |

Note: (1) Xuzhou 142 (upland cotton) with normal fibers was used as a control and compared with two fiber mutants, Xuzhou 142-N without fuzz (fuzz-less), Xuzhou 142-fl without lint and fuzz (lint-less, fuzz-less). (2) The results are average of DNA extracted three times and standard deviation.

